# Chronic Behavioral and Seizure Outcomes following Experimental Traumatic Brain Injury and Comorbid *Klebsiella pneumoniae* Lung Infection in Mice

**DOI:** 10.1101/2024.12.13.628278

**Authors:** Sarah S. J. Rewell, Ali Shad, Lingjun Chen, Erskine Chu, Jiping Wang, Ke Chen, Terence J. O’Brien, Jian Li, Pablo M. Casillas Espinosa, Bridgette D. Semple

## Abstract

Traumatic brain injury (TBI) is a leading cause of long-term disability, and infections such as pneumonia represent a common and serious complication for TBI patients in the acute and subacute post-injury period. While the acute effects of infections have been documented, their long-term consequences on neurological and behavioral recovery as well as the potential precipitation of seizures after TBI remain unclear. This study aimed to investigate the chronic effects of *Klebsiella pneumoniae* infection following TBI, focusing on post-traumatic seizure development and neurobehavioral changes. Using a mouse model, we assessed the long-term effects of TBI and *K. pneumoniae* infection both in isolation and in combination. We found that, while infection with *K. pneumoniae* resulted in loss of body weight and increased mortality compared to vehicle-inoculated mice, there was no additional mortality in TBI animals. Further, although TBI alone induced chronic hyperactivity and reduced anxiety-like behaviors, *K. pneumoniae* lung infection had no lasting effect on these long-term outcomes. Thirdly, while TBI resulted in both spontaneous and evoked seizures long-term post-injury, early post-injury *K. pneumoniae* infection did not affect late onset seizure susceptibility. Together with recent findings on acute outcomes in this combined insult model of TBI and *K. pneumoniae* infection, this study suggests that *K. pneumoniae* does not significantly alter long-term neurobehavioral outcomes or the development of post-traumatic epilepsy. This research highlights the need to further explore the interplay between additional immune insults such as infection that may influence long-term recovery.

## BACKGROUND

Traumatic brain injury (TBI) is a leading cause of long-term disability worldwide, with many survivors suffering from persistent cognitive, emotional, and physical impairments. Among the most significant acute and subacute complications following severe TBI are hospital-acquired infections, particularly pneumonia, which effects a substantial proportion of patients.^1–4^ Pneumonia caused by *Klebsiella pneumoniae*, an opportunistic pathogen commonly associated with ventilator-associated infections, is of particular concern due to its role in exacerbating systemic inflammation and respiratory failure, and increasing the risk of mortality.^4–6^ In TBI patients, immune dysregulation induced by the injury itself heightens vulnerability to such infections, which can worsen functional outcomes and complicate recovery.^7–9^ Several recent studies, including from our group, have investigated the acute consequences of various infection models in the context of TBI and provided important insights into how bacterial infections exacerbate acute post-injury neuroimmune responses.^10–14^ However, there remains a critical gap in understanding how pneumonia following TBI impacts long-term neurological and behavioral outcomes.

Post-traumatic epilepsy (PTE) represents a debilitating long-term complication for survivors of moderate or severe TBI.^15–20^ Defined as recurrent and unprovoked seizures that occur at least one week after TBI,^21^ PTE can severely impact quality of life and is associated with an increased risk of cognitive decline, mood disorders, early-onset neurodegeneration and mortality.^22–26^ Yet the precise mechanisms that drive the development of epileptogenesis after a brain injury remain elusive. Neuroinflammation, a well-known hallmark of TBI, is thought to play a role, and in this way, an additional immune challenge such as an infection is hypothesized to promote post-traumatic epileptogenesis.^27, 28^

To address this hypothesis, we recently conducted a retrospective cohort study examining a large trauma registry of adults with moderate to severe TBI. Infections were documented for approximately one quarter of TBI patients in the registry, with pneumonia being the most common presentation. By multivariate analysis to adjust for known risk factors, we found a solid association between hospital-acquired infections and the development of PTE at 2 years post-injury.^29^ This finding suggests that hospital-acquired infections may contribute to the development of PTE, such that infections represent a modifiable risk factor. However, further exploration of this hypothesis in mouse models have to date failed to produce experimental evidence to support this theory. Specifically, we have evaluated the long-term consequences of peripherally-administered lipopolysaccharide (LPS), as an infection-like immune challenge, after experimental TBI in mice. In both pediatric and adult contexts, we reported that LPS induced a robust acute immune response yet did not exacerbate the long-term development of post-traumatic seizures.^10, 30^

While the LPS mouse model has some advantages as a well-established, predictable model of a systemic immune challenge, it fails to recapitulate many of the key features of a live bacterial infection *in vivo*. As such, it is pertinent that experimental models shift towards preferential use of live infectious agents, delivered via clinically-relevant routes, to more appropriately model the complex pathophysiological scenario of a hospital-acquired infection in an individual with severe TBI.^14^ To address this, we recently established a new model of intratracheal inoculation of *K*. *pneumoniae* bacteria after experimental TBI in the mouse, and conducted detailed characterization of the acute consequences of this dual insult.^11^ We observed that *K. pneumoniae* lung infection after TBI induced a robust yet transient inflammatory response, primarily restricted to the lungs but with some systemic effects, alongside exacerbated elevation of several pro-inflammatory genes such as *Ccl2* in the brain of TBI + *K. pneumoniae* mice. However, the potential long-term consequences of these changes were not determined.

The current study therefore sought to address this knowledge gap by investigating the chronic consequences of *K. pneumoniae* infection following experimental TBI, with a particular focus on the development of post-traumatic seizures and alterations in neurobehavioral outcomes.

## METHODS

### Experimental Timeline

To determine the long-term effects of lung infection with a TBI, mice were subjected to a moderate-to-severe TBI model (or a sham control surgery), followed by intratracheal inoculation with Vehicle or *K. pneumoniae* bacterium at 4 days post-injury. The four experimental groups were Sham-Vehicle, Sham-*Kp*, TBI-Vehicle, and TBI-*Kp*. At approximately 4 months (16 weeks) post-injury, mice underwent extensive behavioral testing over a three-week period. At approximately 4-5 months post-injury, a recording electrode was implanted to allow for subsequent video-EEG monitoring for an average of 10 days per mouse. Finally, all mice received a sub-convulsive dose of pentylenetetrazol (PTZ) to evaluate evoked seizure responses, followed by tissue collection at 6 months post-injury (Figure 1a).

**Figure 1:**
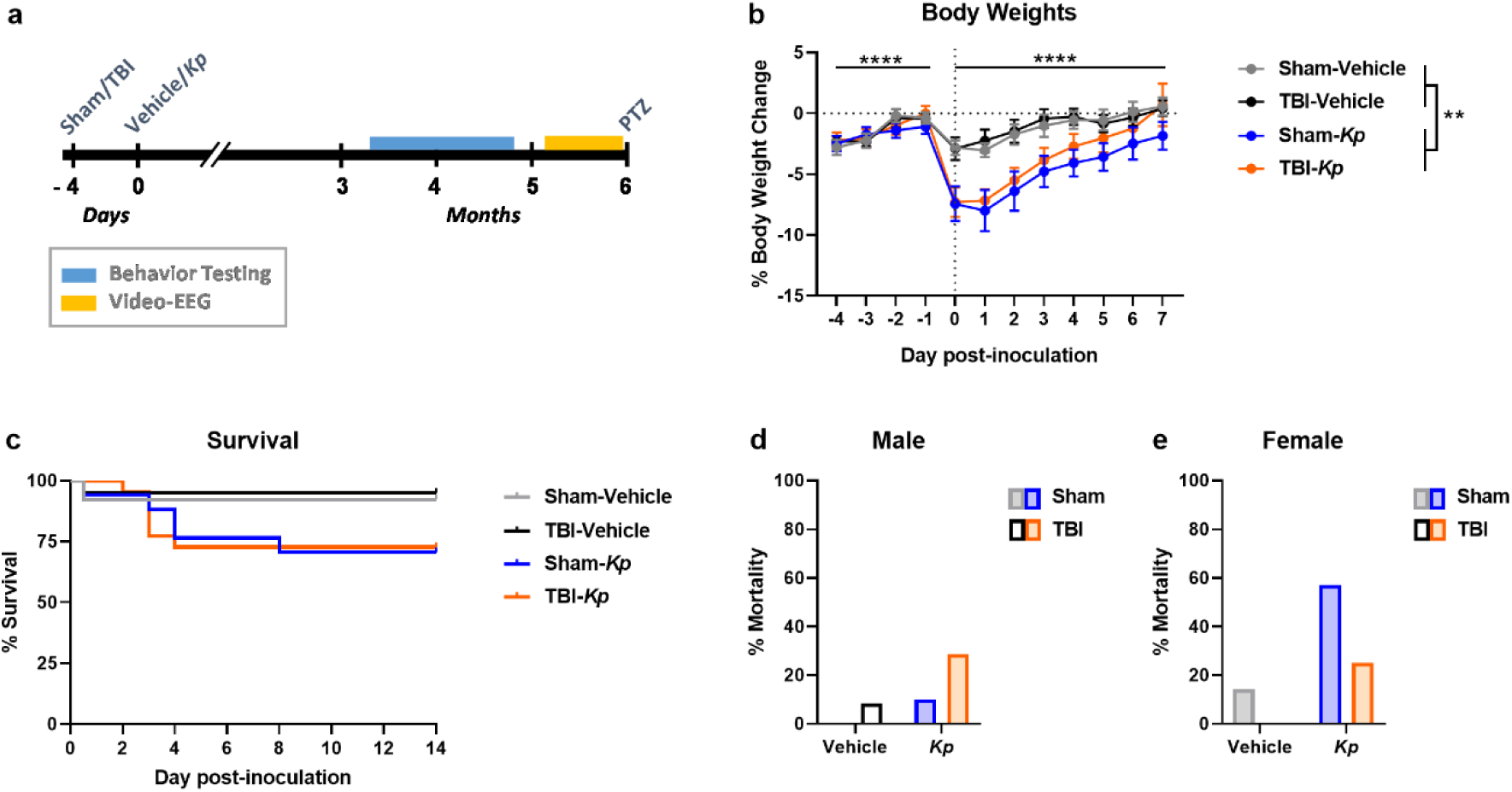
Experimental timeline and acute outcomes. (a) Experimental timeline; (b) Body weight changes from the time of TBI or sham procedure (day -4) then up to 7 days post-inoculation with Kp or Vehicle. **p<0.01 indicates main effect of Kp from a three-way ANOVA. (c) Percent survival over the first two weeks post-inoculation. (d) Mortality rate in males. (e) Mortality rate in females.

### Animals and Ethics

All animal experiments were conducted with approval from the Alfred Research Alliance Animal Ethics Committee (#P8032) and the Animal Care and Use Review Office (ACURO) of the US Department of Defense. These procedures adhered to the approved standards and the Australian Code for the Care and Use of Laboratory Animals as set by the National Health and Medical Research Council of Australia (NHMRC). Male and female C57Bl/6J mice were obtained from the Walter and Eliza Hall Institute of Medical Research in Melbourne, Australia, and acclimated for one week before experiments began. The initial experiments (TBI surgeries and *K. pneumoniae* inoculations) were carried out in QC2 (microbiological containment) facilities at the Monash Research Animal Precinct in Clayton, Australia, before mice were transferred to the Precinct Animal Centre at the Alfred Hospital in Melbourne, for a 4-week quarantine period before behavior testing and video-EEG. The mice were housed in groups of same-sex littermates (2-6 per cage; with mixed experimental conditions per cage) in Optimice® individually-ventilated cages, maintained on a 12-hour light/dark cycle with continuous access to food and water.

### Controlled cortical impact (CCI) model of TBI

Moderate-to-severe experimental traumatic brain injury (TBI) was induced in 10-12 week old mice using the controlled cortical impact (CCI) model, as previously described.^30^ Briefly, anesthesia was initiated with 4% isoflurane in oxygen and maintained at 1.5% via a nose cone. Prior to surgery, all animals were given buprenorphine (0.05 mg/kg in saline; subcutaneously in the flank) and bupivacaine (1 mg/kg in saline; subcutaneously in the scalp) for pain relief, and 0.5 mL of 0.9% saline was administered at the end of the procedure for hydration. The mice were stabilized in a stereotaxic frame, and a 3.5 mm craniotomy was performed over the exposed left parietal bone. An electronic controlled cortical impactor device (eCCE-6.3; Custom Design and Fabrication Inc., Sandston, VA) was used to deliver an impact with a 3 mm rounded tip, at a speed of 4.5 m/s, to a depth of 1.7 mm for 150 ms. Sham animals underwent the same surgical procedure without the impact. After the CCI or sham surgery, the skin incision was sutured and an antiseptic solution applied. The animals were then allowed to recover in individual cages on a heat mat before being returned to their home cage.

### *K. pneumoniae* lung infection model

Freeze-dried cultures of *K. pneumoniae* ATCC 15380 were obtained from In Vitro Technologies (Noble Park, VIC, Australia) and cultured on nutrient agar plates (Medium Preparation Unit, University of Melbourne, Victoria, Australia) at 37 °C overnight. For the initial culture, a large number of *K. pneumoniae* colonies were harvested and suspended in cation-adjusted Mueller-Hinton broth (CAMHB) for an additional 24-hour incubation at 37 °C with shaking at 180 rpm. Mid-logarithmic-phase cultures were then prepared by incubating in fresh CAMHB for 3 hours. The bacterial concentration (colony-forming units (CFU)/mL) was assessed by measuring the optical density (OD) at 600 nm and adjusted to the desired working concentration.

Mice were randomized for inoculation with 10^6^ CFU *K. pneumoniae* or vehicle on day 4 post-injury, as hospital-acquired infections are most common during the first week after a TBI.^31, 32^ A series of pilot studies were previously conducted to determine the optimal dose of *K. pneumoniae* via intratracheal inoculation to achieve an appropriate lung infection model in adult male and female mice.^11^ For inoculation, mice were anesthetized with 2-4% inhaled isoflurane by an experienced technician and a MicroSprayer^®^ Aerosolizer (Penn-Century, Philadelphia, PA, USA) was used to administer 25 μL bacterial suspension or vehicle solution (diluted CAMHB) directly into the trachea.^33^ Five-fold serial dilutions of both *K. pneumoniae* and vehicle samples were spiral plated and incubated overnight at 37 °C on nutrient agar plates to verify the CFU/mL in the administered inoculum, using ProtoCOL 3 software (Synbiosis, USA). Following recovery from the procedure, mice were closely monitored post-treatment for sickness behavior, general appearance, and weight loss over the time course. Weight loss ≥20% was a trigger for immediate humane euthanasia. Animal numbers are depicted in Table 1.

**Table 1:**
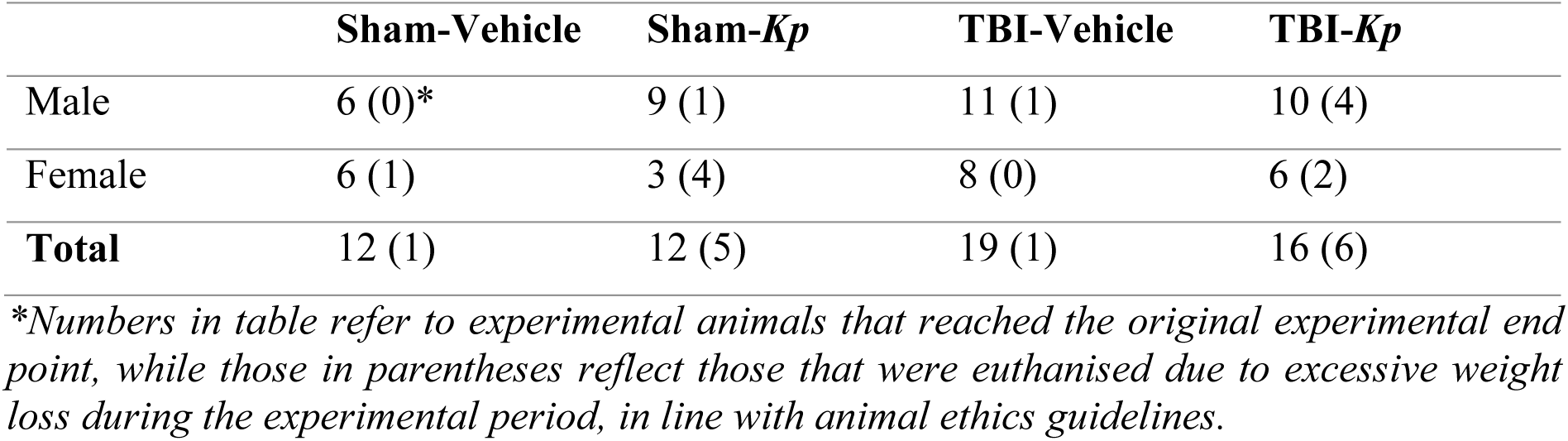
Experimental animal numbers.

|  | Sham-Vehicle | Sham-*Kp* | TBI-Vehicle | TBI-*Kp* |
| --- | --- | --- | --- | --- |
| Male | 6 (0)* | 9 (1) | 11 (1) | 10 (4) |
| Female | 6 (1) | 3 (4) | 8 (0) | 6 (2) |
| **Total** | 12 (1) | 12 (5) | 19 (1) | 16 (6) |

\*Numbers in table refer to experimental animals that reached the original experimental end point, while those in parentheses reflect those that were euthanised due to excessive weight loss during the experimental period, in line with animal ethics guidelines.

### Facility Transfer

The initial experiments (TBI surgeries and *Kp* inoculations) were carried out in QC2 facilities at the Monash Research Animal Precinct in Clayton, Australia, before mice were transferred to the Precinct Animal Centre at the Alfred Hospital in Melbourne for behavior testing and video-EEG monitoring. This experimental design was required due to the specialized facilities required for these procedures. Prior to leaving the Monash Clayton facility, for each of 4 cohorts of mixed-experimental groups, a health screen was performed at 6 weeks post-inoculation by sacrificing and testing a Swiss strain male mouse that was housed alongside the experimental mice and exposed to their bedding weekly. An additional health check was performed at the end of the four-week quarantine period upon arrival at the Alfred Hospital site. This involved the sampling of sera collected from 2-4 experimental animals that had previously had *K. pneumoniae* infection, via submandibular bleed performed by the animal facility technicians. Health screening was performed by Cerberus Sciences (Scoresby, VIC, Australia) via a standard panel. All cohorts passed the health screening. Once through the quarantine period, mice were transferred to the holding rooms in the Department of Neuroscience at Monash University, Alfred Hospital, and habituated for at least one week before behavior testing.

### Behavior Testing

At approximately 16 weeks post-injury, a comprehensive battery of neurobehavioral tests was conducted to assess the chronic consequences of TBI and *K. pneumoniae*. Firstly, mice underwent an Open Field (OF) test, in a square arena for 10 min duration, to evaluate general locomotor activity, exploratory behavior and anxiety-like behavior. Anxiety-like behavior was additionally measured using the Elevated Plus Maze (EPM) during a 10 min period. In both tests, TopScan software was used to track activity and the time spent in the center versus periphery or open arms compared to the closed arms of the maze.^30^

Gross sensorimotor performance was evaluated using the accelerating rotarod test over three consecutive days. Each day, mice completed three trials with a 30-minute rest period between trials. The rotarod device accelerated from 4 to 40 rpm over a 5-minute period, which was the maximum duration of the test. The average latency to fall from the rotarod was calculated for each mouse per day.^30^

Social approach and social novelty preferences were assessed using the three-chamber test, which involved three consecutive 10-minute sessions conducted with a custom-built Perspex apparatus. The test proceeded through three stages: first, a habituation period (stage 1); then, the introduction of a same-sex stimulus mouse into one of the outer chambers (stage 2); and finally, the placement of a second, novel same-sex stimulus mouse into the opposite outer chamber (stage 3). TopScan software was used to monitor the amount of time the experimental mouse spent in each outer chamber. In stage 2, a preference for the chamber with the stimulus mouse over the empty chamber indicated social interest, while in stage 3, a preference for the novel mouse over the familiar mouse demonstrated social recognition or memory.^34, 35^ 2 mice from the TBI-Vehicle group were unable to be analyzed from this test due to aberrant video tracking.

Finally, the sucrose preference test was employed to identify potential depressive-like anhedonia in mice. Over a 5-day period, mice had access to two drinking bottles: one with filtered water and the other with a 1% sucrose solution. The positions of the bottles were switched halfway through the experiment. On the first and last day of the test, the volumes of liquid consumed from each bottle were measured, and a sucrose preference ratio was calculated by dividing the volume of sucrose solution consumed by the total volume of liquid consumed.^10^

### Video Electroencephalography (EEG)

To evaluate chronic seizure activity, EEG electrodes were surgically implanted at 18-19 weeks post-injury. Under isoflurane anesthesia, epidural recording electrodes (E363/20/2.4/SPC ELEC W/SCREW SS, Plastics One Inc., USA) were carefully positioned: one ipsilateral and distal to the craniotomy, one contralateral to the craniotomy (2.5 mm to the right of the midline, -2.5 mm relative to Bregma), and two electrodes over the cerebellum for ground and reference.^36^ An additional anchor screw (00-96 x 3/32, Plastics One Inc., USA) was placed over the left frontal region to reinforce the head cap. All screws were secured to the skull with SuperGlue (Bostik, Australia), and the electrodes were mounted into a pedestal head cap (MS363, Plastics One Inc., USA) and fixed with dental acrylate. The skin was sutured around the head cap, and the animals received subcutaneous pain relief (buprenorphine 0.05 mg/kg and bupivacaine 1 mg/kg) along with saline for hydration. Following electrode implantation, mice were housed individually for the remainder of the experiment.

### Phenotyping Post-Traumatic Epilepsy

From 5 months post-TBI, continuous Video-EEG recordings were collected using a Grael EEG amplifier (Compumedics, Australia), accompanied by infrared video recordings. The EEG data was collected with a high-pass filter at 1 Hz and a low-pass filter at 70 Hz, at a sampling rate of 512 Hz, and digitized using Compumedics Profusion EEG software v 4.0. Recordings were obtained from both ipsilateral and contralateral electrodes, using common ground and reference signals. Each animal had a total of 8-12 days of video-EEG data recorded, which was analyzed using Assyst.^37^ Seizures were identified by changes in the EEG pattern lasting more than 10 seconds, with an amplitude greater than three times the baseline, a repetitive and rhythmic discharge pattern, and variations in amplitude at the start and end of the seizure.^30^ Potential seizure events identified by Assyst were reviewed by an experienced investigator (PMCE), who examined the EEG tracing blinded to the experimental group. Finally, identified seizures were confirmed following review of the video recording. Approximately 20% of video-EEG recordings were of insufficient quality to allow for accurate quantification, and were excluded. The presented results are from the remaining 80% of recordings.

### Pentylenetetrazol Seizure Susceptibility Challenge

Immediately before tissue collection at 24 weeks post-TBI, a single intraperitoneal dose of 40 mg/kg pentylenetetrazol (PTZ) (P6500, Sigma, Australia) was administered to assess susceptibility to evoked seizures as an additional, indirect measure of PTE development. Behavioral responses to PTZ were observed over a 15-minute period. An experienced investigator (SSR), blinded to experimental group, reviewed the video recordings and rated the response to PTZ according to a modified 7-point Seizure Severity Score, where 0 indicates no response or normal activity, and 7 indicates status epilepticus leading to death.^30, 38, 39^

### Postmortem Lesion Assessment

Mice were humanely euthanized via intraperitoneal injection of 160 mg/kg sodium pentobarbitone (Lethabarb®, Virbac, Australia), followed by transcardial perfusion with 4% paraformaldehyde (PFA) at a rate of 2 mL/min. The extracted brains were post-fixed overnight in 4% PFA, then transferred to 70% ethanol and sent to the Monash Histology Platform for paraffin processing and embedding (Monash University, Clayton, Australia). Seven μm coronal brain sections were sectioned and stained with cresyl violet and Luxol Fast Blue, as described previously,^11, 30^ to illustrate the extent of pathology.

### Statistical Analysis

Statistical analysis was performed using GraphPad Prism v.9.4.1 (GraphPad Software Inc., San Diego, CA, USA), with significance defined as p<0.05. Two and three-way analyses of variance (ANOVA) were performed with Tukey’s post hoc test where appropriate. Data with 3 independent variables of time, injury, and infection, were assessed with a 3-way ANOVA. In most instances, both male and female mice are pooled per group, with open circle data points graphically denoting female animals. Potential sex differences were tested by 3-way ANOVA (factors of sex, injury, and infection) where appropriate, and only reported where significant sex differences were detected. Differences in mortality were tested using the Log-Rank (Mantel-Cox) test. Data are presented as mean ± SEM.

## RESULTS

### Impact of *K. pneumoniae* Infection on Body Weight and Mortality in TBI Mice

We sought to test the hypothesis that lung infection with *K. pneumoniae* after a moderate-to-severe experimental TBI would exacerbate chronic behavioral and seizure outcomes. To determine this, four experimental groups (Sham-Vehicle, Sham-*Kp*, TBI-Vehicle, and TBI-*Kp*) were compared across a range of outcome measures (Figure 1a).

Body weights were monitored as an indicator of general health. When compared across the first week post-infection, a reduction in body weight due to *K. pneumoniae* inoculation is evident in Sham-*Kp* and TBI-*Kp* groups (Figure 1b) compared to vehicle-treated groups. Three-way ANOVA confirmed a main effect of time (F_4, 222_=28.36, p<0.0001), a main effect of *Kp* (F_1, 57_=9.36, p=0.0034), and a significant time x *Kp* interaction (F_1, 57_=9.06, p<0.0001). However, no effect of TBI alone was observed (F_1, 57_ = 0.51, p=0.4776).

*K. pneumoniae* inoculation resulted in acute symptoms associated with a lung infection, as expected as described previously in this paradigm.^11^ A portion of mice were euthanized during the first week post-injury/infection, the majority between days 3-5 post-infection, due to excess weight loss ≥20% as per animal ethics guidelines (Table 1). The total mortality rate for vehicle-treated mice was 5.7% (5.3% for males and 6.3% for females), and for *Kp*-infected mice it was 22% (17.2% for males and 27.6% for females). The combined insult of TBI and *Kp* infection appeared to increase mortality in male mice; however, this was not the case for female mice. There was no significant difference between mortality in Sham-*Kp* compared to TBI-*Kp* mice when analyzed over the first two weeks (Figure 1c; Log-Rank (Mantel-Cox) test, Chi-squared =0.0063, p=0.9366).

### Chronic Neurobehavioral Outcomes after TBI and *K. pneumoniae* Infection

At approximately 16 weeks (4 months) post-TBI/Sham and *K. pneumoniae* infection, all experimental mice underwent a battery of neurobehavioral tests to assess long-term functional outcomes. Unless stated, no overt sex differences were observed. In the Open Field test, TBI mice showed an increase in total distance travelled compared to Sham groups (2-way ANOVA, F_1,55_=9.06, p=0.0039; Figure 2a), indicating chronic hyperactivity as previously characterized in this model.^10^ Similarly, TBI mice moved with a higher velocity compared to Sham mice (2-way ANOVA, F_1,55_=11.28, p=0.0014; Figure 2b). However, *Kp* and Vehicle-treated groups performed similarly, and there were not TBI x infection interactions. Further, no effects of either TBI or infection were observed in terms of the proportion of time spent in the center of the arena (p>0.05).

**Figure 2:**
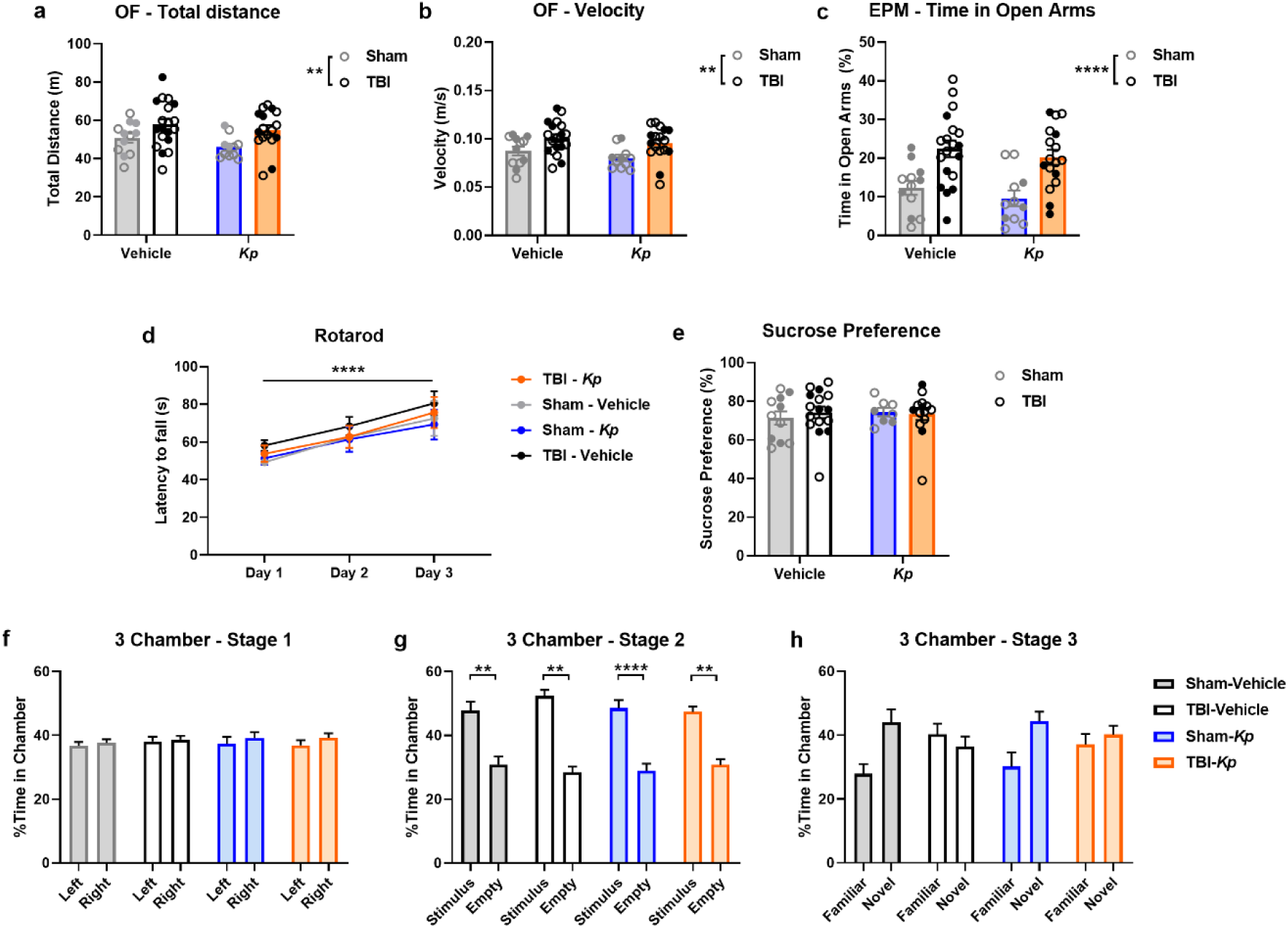
Neurobehavioral outcomes at approximately 4 months post-TBI / Kp infection. (a) Open Field (OF) test, total distance moved. (b) Velocity of movement in the OF test. (c) Time spent in the open arms of the Elevated Plus Maze (EPM). **p<0.01, ****p<0.0001, 2-way ANOVA main effect of TBI. (d) Rotarod, latency to fall. (e) Sucrose preference test. (f) Three-Chamber social approach test, Stage 1; (g) Stage 2; and (h) Stage 3. **p<0.01, ****p<0.0001, 2-way ANOVA main effect of TBI. In Stage 2, **p<0.01, ****p<0.0001 from post-hoc comparisons. Open circles = female; closed circles = male.

Next, the Elevated Plus Maze was used as a measure of anxiety-like behavior (Figure 2c). Here, TBI mice spent significantly more relative time in the open arms of the maze compared to Sham groups (2-way ANOVA, F_1,55_=23.79, p<0.0001), indicating a reduction in anxiety-like behavior chronically after injury as previously described.^10^ Infection did not alter this response. In the Rotarod test (Figure 2d), while all experimental groups demonstrated improvement in performance over time (increased latency to fall from trials 1 to 3), no differences between the groups were observed nor were there any interactions between the factors of TBI, infection or time (3-way RM ANOVA, main effect of time p<0.0001). Similarly, all mice showed a comparable preference for sucrose over water in the Sucrose Preference Test (Figure 2e), of approximately 70-80%, with no effects of TBI or *Kp* (2-way ANOVA, p>0.05).

Finally, the Three-Chamber Social Approach Test was conducted. During habituation in Stage 1 (Figure 2f), as expected, no group differences were observed. In Stage 2 (Figure 2g), a significant main effect of chamber side was detected (3-way ANOVA, F_1,50_=87.71, p<0.0001). Tukey’s post-hoc analyses found that all groups showed a preference for the chamber containing the stimulus animal compared to the empty chamber side, indicating an intact preference for sociability. Lastly, in Stage 3 (Figure 2h), a main effect of chamber side was detected (F_1,50_=6.06, p=0.0173), as well as a chamber side x TBI interaction (F_1,50_=6.63, p=0.0131). Visually, it appears that Sham-Vehicle and Sham-Kp groups spent more time in the chamber containing the novel stimulus mouse compared to the familiar one, while both TBI groups spent roughly equivalent time in each chamber. However, Tukey’s post-hoc analyses failed to detect any specific differences between chamber sides for any of the experimental groups, rendering this stage difficult to interpret.

### Chronic Seizures After TBI and *K. pneumoniae* Infection

Continuous Video-EEG recordings were obtained over an 8-12 day monitoring period per animal at approximately 5 months post-TBI/Sham, to evaluate whether early post-injury *K. pneumoniae* infection altered the development of PTE.

Spontaneous electro-clinical seizure activity was observed in TBI-Vehicle and TBI-*Kp* mice (Figure 3a-b), but none of the sham-operated mice. All seizures were observed to be generalized in nature, commencing almost simultaneously in both hemispheres (ipsilateral and contralateral to the injury) and were typically Racine Class 3 or 4. A total of 7 animals (4 male and 3 female) exhibited at least one spontaneous seizure, and 40% of seizures occurred during the dark phase. All of the TBI-Vehicle mice had one seizure each during the recording period. Of the 3 TBI-*Kp* mice that had seizures, 2 had multiple seizures (Table 2). In total, a comparable proportion of Vehicle and Kp-infected mice were observed to develop PTE: 21.1% of TBI-Vehicle mice (4/19) and 19.7% of TBI-*Kp* mice (3/16) (p>0.999, Fisher’s exact test; Figure 3c). The average seizure duration was 67 sec (median 44 sec; range of 23-238 sec) (Figure 3d). Seizure duration was similar across the two experimental groups (t_8_=0.48, p=0.6421).

**Figure 3:**
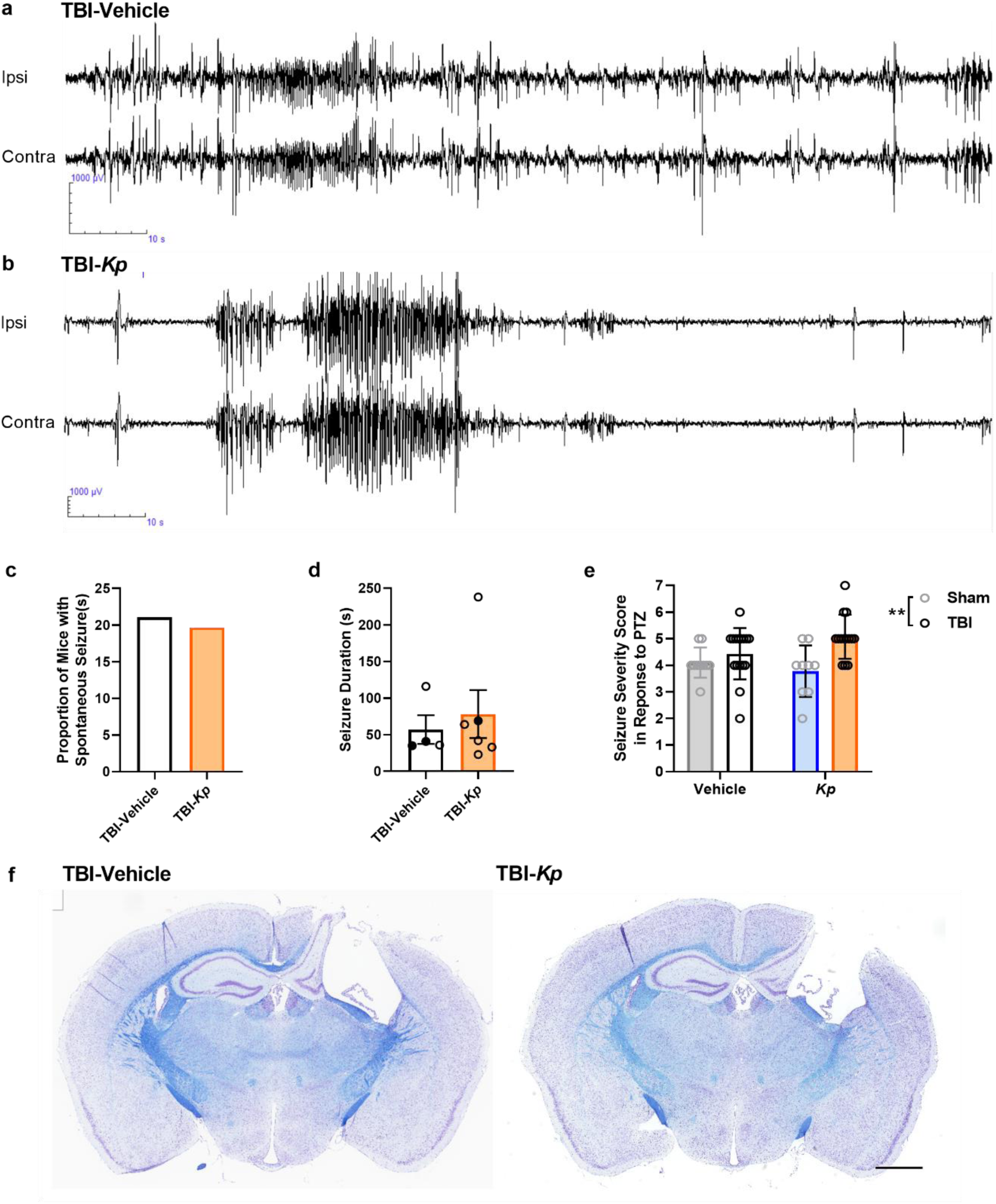
Spontaneous and evoked seizure activity at 5-6 months post-TBI / Kp infection. Representative EEG tracings illustrate spontaneous seizure activity in a TBI-Vehicle (a) and TBI-Kp animal (b) over a 120 sec period. Ipsi = electrode positioned ipsilateral to the injury site; contra = contralateral to the injury site. Quantification (c) found a similar proportion of TBI-Vehicle and TBI-Kp mice exhibiting spontaneous seizures, of a comparable duration (d). Unpaired t-test, p>0.05. Finally, the evoked response to a PTZ challenge was evaluated (e), quantified as the Seizure Severity Score. **p<0.01, 2-way ANOVA main effect of TBI but no effect of Kp. n=12/group for Sham-Vehicle and Sham-Kp; 19 for TBI-Vehicle; 16 for TBI-Kp. Coronal brain sections (f) stained with cresyl violet and Luxol Fast Blue illustrate the typical extent of TBI damage observed following the CCI model at this chronic time point, in both TBI-Vehicle and TBI-Kp mice. Scale bar = 2 mm.

**Table 2:** Spontaneous Seizures Observed During Chronic Video-EEG Monitoring.

| ID | Group | Sex | # Seizures | Racine Class | Duration (sec) | Light or Dark Phase |
| --- | --- | --- | --- | --- | --- | --- |
| 1 | TBI-Vehicle | F | 1 | 4 | 36 | Dark |
| 2 | TBI-Vehicle | M | 1 | 4 | 41 | Dark |
| 3 | TBI-Vehicle | F | 1 | 3 | 116 | Light |
| 4 | TBI-Vehicle | M | 1 | 4 | 35 | Light |
| 5 | TBI-Kp | M | 1 | 3 | 69 | Dark |
| 6 | TBI-Kp | F | 2 | 4 | 33 | Dark |
|  |  |  |  | 4 | 42 | Light |
| 7 | TBI-Kp | F | 3 | 3 | 238 | Light |
|  |  |  |  | 4 | 64 | Light |
|  |  |  |  | 1 | 23 | Light |

Upon completion of continuous video-EEG recording, mice were administered 40 mg/kg i.p. PTZ and observed for a 15 min period. The behavioral response to PTZ was video-recorded then scored by a blinded investigator according to the modified Seizure Severity Score^10, 36^ (Figure 3e). 2-way ANOVA revealed that the average Seizure Severity Score in response to PTZ was higher in TBI compared to Sham mice (main effect of TBI, F_1,45_ = 10.41, p=0.0023), while *Kp* had no effect (F_1, 45_ = 0.38, p=0.5406).

All mice were euthanized after the PTZ challenge and brain tissue collected for histology. A representative image of a TBI-Vehicle brain is depicted in Figure 3f, illustrating the typical pattern of unilateral cortical and hippocampal damage in the chronic phase as a result of the CCI model.

## DISCUSSION

*K. pneumoniae* is a leading cause of ventilator-acquired pneumonia in critically-ill and immunocompromised individuals, and is common in patients with severe TBI.^31, 40–43^ After recently establishing a new mouse model of experimental TBI combined with a pulmonary *K. pneumoniae* infection,^11^ in the current study we sought to investigate if the resolved infection had chronic consequences for TBI outcomes.

In previous pilot studies, we determined 1 x 10^6^ CFU of *K. pneumoniae* ATCC 15380 to be most appropriate for experimental use.^11^ Here, we found that this dose induced considerable body weight and either spontaneous mortality or substantial weight loss triggering humane euthanasia for approximate 20% of infected mice (∼15% higher than mortality for Vehicle-treated mice). All mortality occurred within several days of infection, presumably due to pathology associated with the robust innate immune response triggered by *K. pneumoniae* inoculation, as demonstrated previously.^11^ However, the combined insult of TBI plus infection did not exacerbate body weight loss, nor was there additional mortality observed in the TBI-*Kp* group compared to Sham-*Kp* mice. Of note, however, sex differences in mortality were observed. While females overall appeared to be more susceptible to *K. pneumoniae* infection, the combination of TBI+*Kp* induced the highest mortality rate in male mice. With mixed reports of sex-specific responses to *K. pneumoniae* from experimental models,^11, 44–46^ potential sex-specific responses to infection warrant further investigation.

Turning to long-term outcomes, we focused on neurobehavioral function and post-traumatic epilepsy. Of note, we have previously demonstrated in this model that most of the *K. pneumoniae* bacteria have been cleared from the lungs of surviving mice by 7 days post-infection, and certainly by a 28 day time point.^11^ Here, we further confirmed that mice had resolved their infection prior to transfer between facilities for behavior testing and video-EEG, and no differences were observed between body weight trajectories of any experimental group after the initial two weeks (data not shown).

We considered it plausible that prior K. pneumoniae infection, despite resolution, may nonetheless have long-term consequences for neurological function. For example, long-term deficits in exploratory locomotor behavior were reported in a mouse model of intranasal *K. pneumoniae*-induced pneumosepsis, associated with persistent brain inflammatory gene expression.^47^ We have previously demonstrated that lung infection with *K. pneumoniae* alone induced acute expression of *Tnfα* in the cortex; while the combination of TBI plus *K. pneumoniae* resulted in elevated brain gene expression of pro-inflammatory cytokine *Ccl2* and oxidative stress mediator *Hmox1*.^11^ However, at 4 months post-injury, we found that previously-infected *K. pneumoniae* mice were indistinguishable from Vehicle-treated mice across a range of tests for exploratory activity, sensorimotor function, anxiety-like behavior, social cognition, and pleasure-seeking behavior. TBI mice displayed chronic hyperactivity and reduced anxiety-like behavior as expected in this model of CCI.^10, 30^

The primary outcome of this study was the evaluation of post-traumatic seizures, based on recent epidemiological evidence of an association between hospital-acquired infection and the development of PTE by 2 years post-injury in patients.^48^ From extended continuous video-EEG monitoring, we observed spontaneous seizure activity in approximately 20% of TBI mice, regardless of prior *K. pneumoniae* infection, and their seizures were of a similar nature and duration. In addition, both TBI groups showed a similar response to the PTZ challenge, an indirect measure of seizure susceptibility indicating epileptogenesis. This finding is in line with our previous experimental work using a similar experimental design but LPS as a mimic infectious agent, where LPS failed to have lasting effects on either neurobehavioral or seizure outcomes.^10^

Together, these studies now suggested that either (a) there is not a robust or direct relationship between infection and these chronic outcomes; or (b) that the experimental models are still inadequate to replicate the clinical scenario. Of note, while live *K. pneumoniae* in this study was administered into the lungs (a clinically-relevant route), it was delivered as a single bolus which does not represent how a bacterial load expands over a time course in infected patients.^14, 49, 50^ This may account for some of the mortality we observed, but may also influence the resulting immune response including brain-lung-immune interactions. Further, inherent differences between human and rodent immunity may render the mouse an insufficient model system for investigating the complex scenario of a critically-ill patient with TBI and infection.^14^

Several strengths and limitations of the current study are worth noting. The study was carefully designed to dissect out specific effects of the combined insult of TBI plus *K. pneumoniae* compared to either TBI or infection alone, and is one of few studies to date to consider the long-term consequences of such a paradigm. Detailed seizure monitoring via continuous video-EEG is the gold-standard measure of PTE, and inclusion of both male and female mice allowed for consideration of typical biological variability as well as exploration of potential sex differences in responses to injury and infection. Conversely, the study was limited by lack of longitudinal analysis of seizure development, which could provide additional insight into seizure onset and progression over time. Use of a single infection model, time of administration relative to injury, and host strain/species, also limits the generalizability of the findings to other types of infections or pathogens. Finally, while this study focused on behavioral and seizure outcomes, we did not delve into the biological or molecular mechanisms that underlie these presentations. Future studies may benefit from the complementary inclusion of detailed histological or molecular analyses.

## CONCLUSION

In conclusion, this study provides important insights into the potential long-term effects of *K. pneumoniae* infection following moderate-to-severe TBI. We found that a non-trivial *K. pneumoniae* lung infection, associated with ∼20% mortality, did not affect the chronic TBI-induced neurobehavioral deficits or PTE for the survivors. These findings suggest that while *K. pneumoniae* can worsen acute health parameters, its impact on long-term neurological outcomes post-TBI is limited once the infection has been resolved. Further research is needed to further explore potential sex-specific responses to infection, and consider how the timing of infection relative to injury may influence the host’s response. Overall, this work contributes to our growing understanding of how infections influence brain injury outcomes. Given the high incidence of both infections and seizures in TBI patients, and a growing concern over multidrug-resistant pathogens, such research is vital to the medical management of infections in TBI patients and support of optimal long-term outcomes.

## Acknowledgements

The authors would like to thank the Monash Animal Research Platform (Clayton and Alfred Precincts) for their assistance with the project.

## Funding

This project was supported by an Epilepsy Research Program Idea Development Award (#W81XWH-19-ERP-IDA) from the US Department of Defense, awarded to BDS, JL and TOB. BDS was also supported by a Veski Near-Miss Grant. TOB is supported by a National Health and Medical Research Council of Australia (NHMRC) Investigator Grant (APP1176426). JL is supported by an NHMRC Principal Research Fellowship (APP1157909). PMCE is supported by the NHMRC (APP1087172 and APP2013629), a Monash Future Leader Fellowship (FLPF24-0237761657), a Medical Research Future Fund (MRFF) Stem Cell Therapy Missions Grant (MRF1201781), and the US Department of Defense USA Epilepsy Research Program (DoD ERP IDA, grant # EP200022, DoD ERPA RPA, grant# EP220067).

## Author Contributions

Project conceptualization: BDS, TOB, JL. Data access and analysis: BDS, AS, SSR, LC, EC, KC, JW, PMCE. Manuscript draft: BDS. Manuscript edits: All authors. We confirm that this manuscript is consistent with the Journal’s and publisher’s position on issues involved in ethical publication.

## Ethics Approval

All animal experiments were conducted following approval from the local Alfred Research Alliance Animal Ethics Committee (#P8032) as well as the Animal Care and Use Review Office (ACURO) from the Office of Research Protections, US Department of Defense, and carried out in accordance with these approved standards as well as the Australian Code for the Care and Use of Laboratory Animals as stipulated by the National Health and Medical Research Council of Australia (NHMRC).

## Conflicts of Interest

The authors do not have any conflicts of interest to declare. The funders had no role in the design of the study; in the collection, analyses, or interpretation of data; in the writing of the manuscript; or in the decision to publish the results.

## Data Availability Statement

The data that support the findings of this study are available from the corresponding author upon reasonable request.

## Notes

### Competing Interest Statement

The authors have declared no competing interest.

